# Out with the old and in with the new: The role of intolerance of uncertainty in reversal of threat and safety

**DOI:** 10.1101/386797

**Authors:** Jayne Morriss, Francesco Saldarini, Catherine Chapman, Miriam Pollard, Carien M. van Reekum

**Affiliations:** Centre for Integrative Neuroscience and Neurodynamics, School of Psychology and Clinical Language Sciences, University of Reading Reading, UK

**Keywords:** Threat, Conditioning, Acquisition, Reversal, Uncertainty, Anxiety

## Abstract

The ability to learn and reverse threat associations is crucial for survival. The extent to which old threat associations are inhibited and new threat associations are formed may depend on sensitivity to future threat uncertainty. To assess the extent to which Intolerance of Uncertainty (IU) predicts threat learning and reversal, we recorded expectancy ratings and skin conductance in 44 healthy participants during an associative learning paradigm, where threat and safety contingencies were reversed. During acquisition and reversal, we observed larger SCR magnitude and expectancy ratings for threat vs. safety cues. However, during reversal higher IU was associated with larger SCR magnitude to new threat vs. new safety cues, compared to lower IU. These results were specific to IU-related variance, over shared variance with trait anxiety (STAIX-2). Overall, these findings suggest that individuals high in IU are able to reverse threat and safety associations in the presence of direct threat. Such findings help us understand the recently revealed link between IU and threat extinction, where direct threat is absent. Moreover, these findings highlight the potential relevance of IU in clinical intervention and treatment for anxiety disorders.

## 1. Introduction

The ability to discriminate and adjust behaviour to stimuli that predict threatening or safe outcomes is vital for survival and protection against anxiety and stress disorders (LeDoux, 1998; Milad & Quirk, 2012; Shin & Liberzon, 2009). An organism can learn to associate neutral cues (conditioned stimulus, e.g. a visual stimulus such as a shape) with threatening outcomes (unconditioned stimulus, e.g. shock, loud tone) or safe outcomes. Repeated presentations of a neutral cue with a threatening outcome then results in defensive responding to the cue alone (conditioned response). Importantly, learned threat and safe associations can be updated when the outcome changes.

In animal studies, partial reinforcement, particularly 50%, has been shown to maintain the conditioned response during threat extinction (Jenkins & Stanley Jr, 1950; Leonard, 1975; Livneh & Paz, 2012). However, only a few studies in humans have directly compared different partial reinforcement rates (Chin, Nelson, Jackson, & Hajcak, 2016; Dunsmoor, Bandettini, & Knight, 2007), and shown slower threat extinction after partial reinforcement, compared to continuous reinforcement (Grady, Bowen, Hyde, Totsch, & Knight, 2016; Leonard, 1975). Under partial reinforcement, it is thought that the conditioned response is maintained due to uncertainty regarding the threatening outcome.

Uncertainty has recently been identified as an important factor in anxiety and stress disorders (Carleton, 2016a, 2016b; Dugas, Buhr, & Ladouceur, 2004; Grupe & Nitschke, 2013). Notably, recent research from our lab and others has shown that individual differences in Intolerance of Uncertainty (IU), a dispositional tendency to find uncertain situations aversive and anxiety provoking (Carleton, 2016a, 2016b; Dugas, Buhr, & Ladouceur, 2004), plays a critical role in threat acquisition (Chin, Nelson, Jackson, & Hajcak, 2016) and extinction (Dunsmoor, Campese, Ceceli, LeDoux, & Phelps, 2015; Morriss, Christakou, & Van Reekum, 2015, 2016; Morriss, Macdonald, & van Reekum, 2016). Effects of IU on threat acquisition have been found using a variety of measures such as skin conductance (Morriss, Macdonald, & van Reekum, 2016), startle (Chin, Nelson, Jackson, & Hajcak, 2016; Sjouwerman, Scharfenort, & Lonsdorf, 2017) and ratings (Morriss, Macdonald, & van Reekum, 2016; Sjouwerman, Scharfenort, & Lonsdorf, 2017). Effects of IU on threat extinction have been found primarily with skin conductance (Dunsmoor, Campese, Ceceli, LeDoux, & Phelps, 2015; Morriss, Christakou, & Van Reekum, 2015, 2016; Morriss, Macdonald, & van Reekum, 2016), and brain imaging (Morriss, Christakou, & Van Reekum, 2015), and have less consistently been observed in ratings. These results suggest that there may be overlap between different read-out measures for the effect of IU on threat acquisition and extinction.

However, there remain some nuances for threat acquisition and IU. For example, under 50% reinforcement during acquisition, high IU individuals have been shown to exhibit greater discrimination in startle response – indicative of a negative affective background state - to threat versus safety cues (Chin, Nelson, Jackson, & Hajcak, 2016). Furthermore, during an acquisition phase with 50% reinforcement and generalization stimuli (cues that look similar to the CS), high IU individuals have also been observed to show greater generalization in skin conductance response – a measure of sympathetic arousal - across threat and safety cues (Morriss, Macdonald, & van Reekum, 2016). Findings are more consistent for threat extinction (contingencies are updated from threat to safe) after continuous or partial reinforcement, such that high IU individuals have been found to show generalised skin conductance response across threat and safety cues during early extinction, and to show continued skin conductance responding to threat versus safety cues during late extinction (Morriss, Christakou, & Van Reekum, 2015, 2016; Morriss, Macdonald, & van Reekum, 2016). Despite the differences in findings for different read-out measures, overall the results from the literature suggest that IU may act as an important modulator of associative learning mechanisms.

When updating threat to safe associations during extinction, there is a period of uncertainty regarding the change of outcome, and this may induce uncertainty-related anxiety in high IU individuals. However, it is unknown whether high IU individuals have difficulty with: (1) generally updating threat to safe associations when contingencies change, or (2) specifically with updating threat to safe associations during extinction where the US omitted. This question can be examined by adopting a threat reversal paradigm, where both threat and safety contingencies are reversed (Costa, Bradley, & Lang, 2015; Kluge et al., 2011; Li, Schiller, Schoenbaum, Phelps, & Daw, 2011; Mertens & De Houwer, 2016; Morris & Dolan, 2004). Through reversal learning, the flexibility of threat and safe associations can be assessed.

Despite the richness that threat reversal paradigms offer in understanding the flexibility of learned associations, little research has been conducted on threat reversal in relation to individual differences in anxiety. Given the important role of uncertainty in anxiety (Carleton, 2016a, 2016b; Grupe & Nitschke, 2013) and that current therapies are based on associative learning principles (Milad & Quirk, 2012), examining the link between IU and the reversal of threat and safe associations may provide crucial information relevant to anxiety disorder pathology and treatment.

Here we used threat and safety cues during acquisition and reversal, in order to assess the relationship between individual differences in self-reported IU and updating of learned threat and safety associations. We measured skin conductance response (SCR) and expectancy ratings whilst participants performed the acquisition and reversal phases. We used an aversive sound as an unconditioned stimulus and visual shape stimuli as conditioned stimuli, similar to previous conditioning research (Morriss, Christakou, & Van Reekum, 2015, 2016; Morriss, Macdonald, & van Reekum, 2016; Neumann, Waters, & Westbury, 2008; Phelps, Delgado, Nearing, & LeDoux, 2004). We used a 50% reinforcement rate during both acquisition and reversal to maintain conditioning (Jenkins & Stanley Jr, 1950) and induce greater uncertainty.

In general, we hypothesised that, during threat acquisition and reversal, skin conductance responding and expectancy ratings would be higher to the threat (CS+) versus safety cues (CS-). Furthermore, we had two alternative hypotheses for IU and updating of learned threat and safety associations during acquisition. Based on previous opposing findings in the literature, during acquisition high IU individuals would be prone to either (1) greater discrimination (Chin et al., 2016) indicated by larger SCR and expectancy ratings to the threat versus safety cues, or (2) less discrimination, and have larger SCR and expectancy ratings for both threat and safety cues (Morriss, Macdonald, et al., 2016). Given the lack of research on reversal and IU we based our hypotheses on the extinction and IU literature. If high IU individuals generally have difficulty updating threat to safe associations when contingencies change then we should observe less discrimination in SCR and expectancy ratings for threat and safety cues during reversal. However, if high IU individuals only have difficulty when updating threat and safety associations during extinction when the US is omitted, we should observe reversal of threat and safety associations for the SCR and ratings. For both acquisition and reversal, we tested the specificity of IU effects by controlling for individual variation reported on the commonly used Spielberger State-Trait Anxiety Inventory, Trait Version (STAIX-2) (Spielberger, Gorsuch, Lushene, Vagg, & Jacobs, 1983).

## 2. Method

### 2.1 Participants

44 students took part in this study (*M* age = 20.45, *SD* age = 3.18; 33 females & 11 males). The sample size is within the same range of previous experiments using psychophysiological measures to examine conditioning and individual differences in anxiety (e.g. Chin, Nelson, Jackson, & Hajcak, 2016; Morriss, Christakou, & Van Reekum, 2016). For this study, the sample size was not based on a formal power calculation. All participants had normal or corrected to normal vision. Participants were recruited through the University of Reading Psychology Panel. The procedure was approved by the University of Reading’s Research Ethics Committee.

### 2.2 Procedure

Participants arrived at the laboratory and were informed on the procedures of the experiment. Firstly, participants were taken to the testing booth and given a consent form to sign as an agreement to take part in the study. Secondly, to assess anxious disposition we asked participants to complete a series of questionnaires presented on a computer in the testing booth. Next, physiological sensors were attached to the participants’ non-dominant hand. Participants were simply instructed to: (1) maintain attention to the task by looking and listening to the coloured squares and sounds presented, (2) respond to the rating scale that followed each block (see “Conditioning task” below for details) using the keyboard with their dominant hand and (3) to sit as still as possible. Participants were presented a conditioning task on the computer, whilst electrodermal activity, interbeat interval and ratings were recorded. Altogether, the experiment took approx. 25 minutes.

### 2.3 Conditioning task

The conditioning task was designed using E-Prime 2.0 software (Psychology Software Tools Ltd, Pittsburgh, PA). Visual stimuli (CS+ and CS-) were presented using a screen resolution of 800 × 600 with a 60 Hertz refresh rate. Participants sat at approximately 60 cm from the screen. Visual stimuli were light blue and yellow squares with 183 × 183 pixel dimensions that resulted in a visual angle of 5.78º × 9.73°. The sound stimulus (US) consisted of a fear inducing female scream (Morriss, Christakou, & Van Reekum, 2015, 2016; Morriss, Macdonald, & van Reekum, 2016).

Acquisition and reversal phases were each presented in two separate blocks, totaling four blocks for the experiment as a whole (see Fig. 1). In acquisition, one of the squares (blue or yellow) was paired with the aversive sound 50% of the time (CS+), whilst the other square (yellow or blue) was presented alone (CS−). The 50% pairing ratio was designed to maximise the unpredictability of the US following the CS+. In reversal, the CS+ and US association was reversed so that the former CS+ became the new CS- and the former CS- became the new CS+. The 50% pairing ratio was maintained during the reversal phase. Participants were naïve to the conditioning procedure, the pairing rate, and contingency change during reversal; all they were told in-between phase is that they could take a break and were asked whether they agreed and were ready to continue. The acquisition phase consisted of 24 trials (6 CS+ paired, 6 CS+ unpaired, 12 CS−) and the reversal phase consisted of 32 trials (8 new CS+ paired, 8 new CS+ unpaired, 16 CS−). Each block therefore consisted of 12 trials during acquisition, and 16 trials during reversal. The reversal phase was longer to allow time for learning to change. Experimental trials within the conditioning task were pseudo-randomised: The first trial of the acquisition and reversal phases started with a trial that was paired. Thereafter the order of all remaining trials were fully randomised. Conditioning contingencies were counterbalanced, with half of the participants receiving the US with a blue square and the other half of participants receiving the US with a yellow square. The presentation times of the task were: 4000 ms square, 1000 ms sound (co-terminated with the square), 6000 - 8800 ms intertrial interval (see Fig. 1).

**Fig 1.**
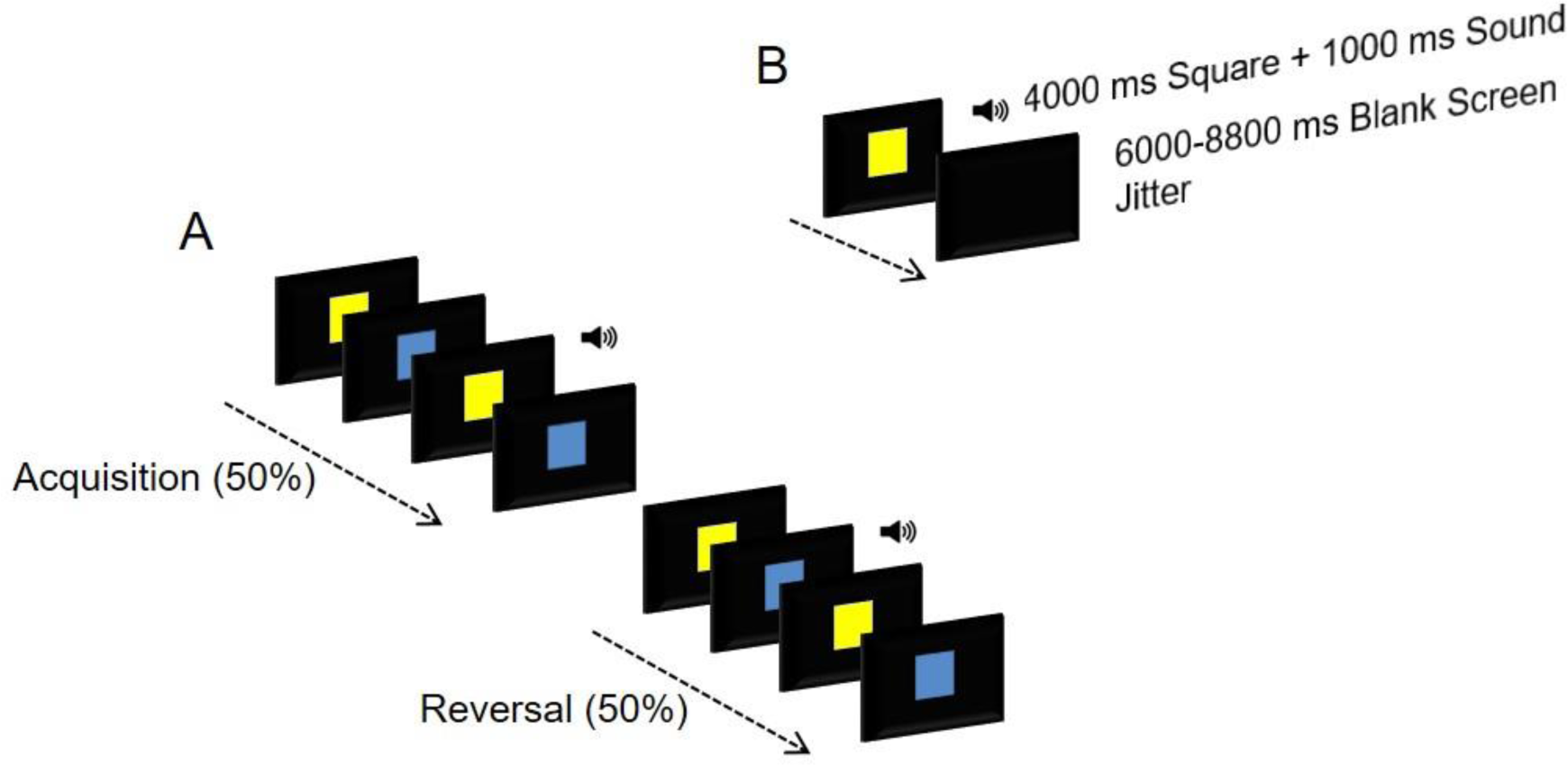
Image depicting (A) the experimental phases of the experiment, and (B) examples of trial timing.

At the end of each block participants were asked to rate the expectancy of the sound stimulus when preceded by the blue square or the yellow square using 9-point Likert scales ranging from 1 ("Don’t Expect") to 9 ("Do expect"). Two other 9-point Likert scales were presented at the end of the experiment. The first one asked participants to rate the valence of the sound stimulus from 1 (negative) to 9 (positive). The second one asked the participants to rate the arousal of the sound stimulus from 1 (calm) to 9 (excited).

### 2.4 Questionnaires

To assess anxious disposition, we presented the following questionnaires on a computer: Spielberger’s State-Trait Anxiety Inventory, Trait Version (STAIX-2) (Spielberger et al., 1983) and Intolerance of Uncertainty (IU) (Buhr & Dugas, 2002). Similar distributions and internal reliability of scores were found for the anxiety measures, STAIX-2 (*M* = 43.11; *SD* = 10.85; range = 25-67; *α* = .93), IU (*M* = 63.3; *SD* = 22.3; range = 28-111; *α* = .96). The STAIX-2 and IU scores were significantly positively correlated, *r*(42) =.81, *p* < .001.

### 2.5 Rating data scoring

Rating data were reduced for each subject by calculating their average responses for each experimental condition using the E-Data Aid tool in E-Prime (Psychology Software Tools Ltd, Pittsburgh, PA).

### 2.6 Physiological acquisition and scoring

Physiological recordings were obtained using AD Instruments (AD Instruments Ltd, Chalgrove, Oxfordshire) hardware and software. Electrodermal activity was measured with dry MLT116F silver/silver chloride bipolar finger electrodes that were attached to the distal phalanges of the index and middle fingers of the non-dominant hand. A low constant-voltage AC excitation of 22mVrms at 75 Hz was passed through the electrodes, which were connected to a ML116 GSR Amp, and converted to DC before being digitized and stored. Interbeat Interval (IBI) was measured using a MLT1010 Electric Pulse Transducer, which was connected to the participant’s distal phalange of the ring finger. An ML138 Bio Amp connected to an ML870 PowerLab Unit Model 8/30 amplified the electrodermal and interbeat interval signals, which were digitized through a 16-bit A/D converter at 1000 Hz. IBI signal was used only to identify movement artefacts and was not analyzed. The electrodermal signal was converted from volts to microSiemens using AD Instruments software (AD Instruments Ltd, Chalgrove, Oxfordshire).

CS+ paired trials were discarded from the analysis to avoid the sound confound (for examination of US responses to the CS+paired trials, see supplementary material). Data from the CS+ unpaired and CS- trials were included. Skin conductance responses (SCR) were scored when there was an increase of skin conductance level exceeding 0.03 microSiemens. The amplitude of each SCR was scored as the difference between the onset and the maximum deflection prior to the signal flattening out or decreasing. SCR onsets and respective peaks were counted if the SCR onset was within 0.5-3.5 seconds (CS response) following CS onset (Dunsmoor, Murty, Davachi, & Phelps, 2015; Hartley, Fischl, & Phelps, 2011; Spoormaker et al., 2011). Trials with no discernible SCRs were scored as zero.

SCR amplitudes were square root transformed to reduce skewness (Dawson, Schell, & Filion, 2000). Trials with motion artefacts, as identified by distortions in both electrodermal and IBI signals, were discarded from the analysis. 0.2 % (4 out of 1,848) of trials were removed from the analysis due to movement artefacts. SCR magnitudes were calculated from remaining trials by averaging SCR square root transformed values and zeros for each contingency and block, creating the following conditions: Acquisition CS+ Early; Acquisition CS- Early; Acquisition CS+ Late; Acquisition CS- Late; Reversal New CS+ Early; Reversal New CS- Early; Reversal New CS+ Late; Reversal New CS- Late. SCR magnitudes were finally z-scored to control for interindividual differences in skin conductance responsiveness (Ben-Shakhar, 1985) related to dryness and thickness of the skin and the surface area of the finger relative to the skin conductance electrode.

### 2.7 Learning assessment

To assess whether participants learned the association between the neutral cue and aversive sound, we calculated separate conditioned response scores for ratings and SCR magnitude from the acquisition and reversal phases. The conditioned response scores were the CS+ trials – the CS- trials for each phase. A positive differential response score indicated a larger response for CS+ relative to CS-, indexing a conditioned response. The learning criterion procedure has been suggested to be problematic (Lonsdorf et al., 2017), however, we used it for comparison with other laboratories that commonly use this procedure. We considered participants ‘learners’ if they displayed a positive differential response in either phase (Ratings: Acquisition 43 learners, 1 non-learner; Reversal 41 learners, 3 non-learners; SCR: Acquisition 34 learners, 10 non-learners; Reversal 25 learners, 19 non-learners). A similar learning criterion has been published elsewhere (Morriss, Macdonald, & van Reekum, 2016). Based on this criterion, only four participants out of the forty-four participants displayed no differential response in both acquisition and reversal. However, as removing these participants did not change the results reported here, for reasons of completeness we decided to include these four participants.

### 2.8 Ratings and SCR magnitude analysis

The analysis was conducted using the mixed procedure in SPSS 21.0 (SPSS, Inc; Chicago, Illinois). We conducted separate multilevel models on ratings and SCR magnitude from acquisition and reversal. For ratings and SCR magnitude we entered Stimulus (CS+, CS-), and Time (Early, Late) at level 1 and individual subjects at level 2. We included the following individual difference predictor variables into the multilevel models: IU, STAIX-2. In all models, we used a diagonal covariance matrix for level 1. Random effects included a random intercept for each individual subject, where a variance components covariance structure was used. Fixed effects included Stimulus and Time. We used a maximum likelihood estimator for the multilevel models.

Where a significant interaction was observed with IU, we performed follow-up pairwise comparisons on the estimated marginal means of the relevant conditions estimated at specific IU values of + or −1 SD of mean IU, adjusted for the control variable (STAIX-2). These data are estimated from the multilevel model of the entire sample, not unlike performing a simple slopes analysis in a multiple regression analysis. Similar analyses have been published elsewhere (Morriss, Macdonald, & van Reekum, 2016; Morriss, McSorley, & van Reekum, 2017).

## 3. Results

### 3.1 Ratings

On average, the participants rated the sound stimulus as aversive (M = 2.05 SD = .91, range 1-4, where 1 = very negative and 9 = very positive) and arousing (M = 7.11, SD = 1.33, range 4-9 where 1 = calm and 9 = excited).

For the expectancy ratings, during acquisition participants reported greater expectancy of the sound with the CS+, compared to CS- [Stimulus: *F*(1, 123.240) = 659.340, *p* < .001] (for descriptive statistics of ratings see Table 1 and Fig. 2a).

**Table 1.** Summary of means (SD) for each dependent measure as a function of condition during the acquisition and reversal phase

| Measure | Acquisition |  |  |  | Reversal |  |  |  |
| --- | --- | --- | --- | --- | --- | --- | --- | --- |
|  | Early |  | Late |  | Early |  | Late |  |
|  | CS+ | CS- | CS+ | CS- | CS+ | CS- | CS+ | CS- |
| SCR |  |  |  |  |  |  |  |  |
| magnitude | 0.55 | 0.09 | 0.31 | -0.15 | 0.04 | -0.13 | -0.02 | -0.16 |
| ($\sqrt{\mu\text{S}}$) | (0.79) | (0.48) | (0.61) | (0.41) | (0.67) | (0.28) | (0.56) | (0.29) |
| Expectancy | 6.23 | 1.64 | 6.89 | 1.27 | 6.50 | 3.23 | 6.70 | 2.55 |
| rating | (1.74) | (1.43) | (1.48) | (0.59) | (1.34) | (1.80) | (1.62) | (1.80) |

Note: Expectancy rating, 1 = "Don't Expect" and 9 = "Do Expect"; SCR magnitude ( $\sqrt{\mu\text{S}}$ ), square root transformed and z-scored skin conductance magnitude measured in microSiemens

**Fig 2.**
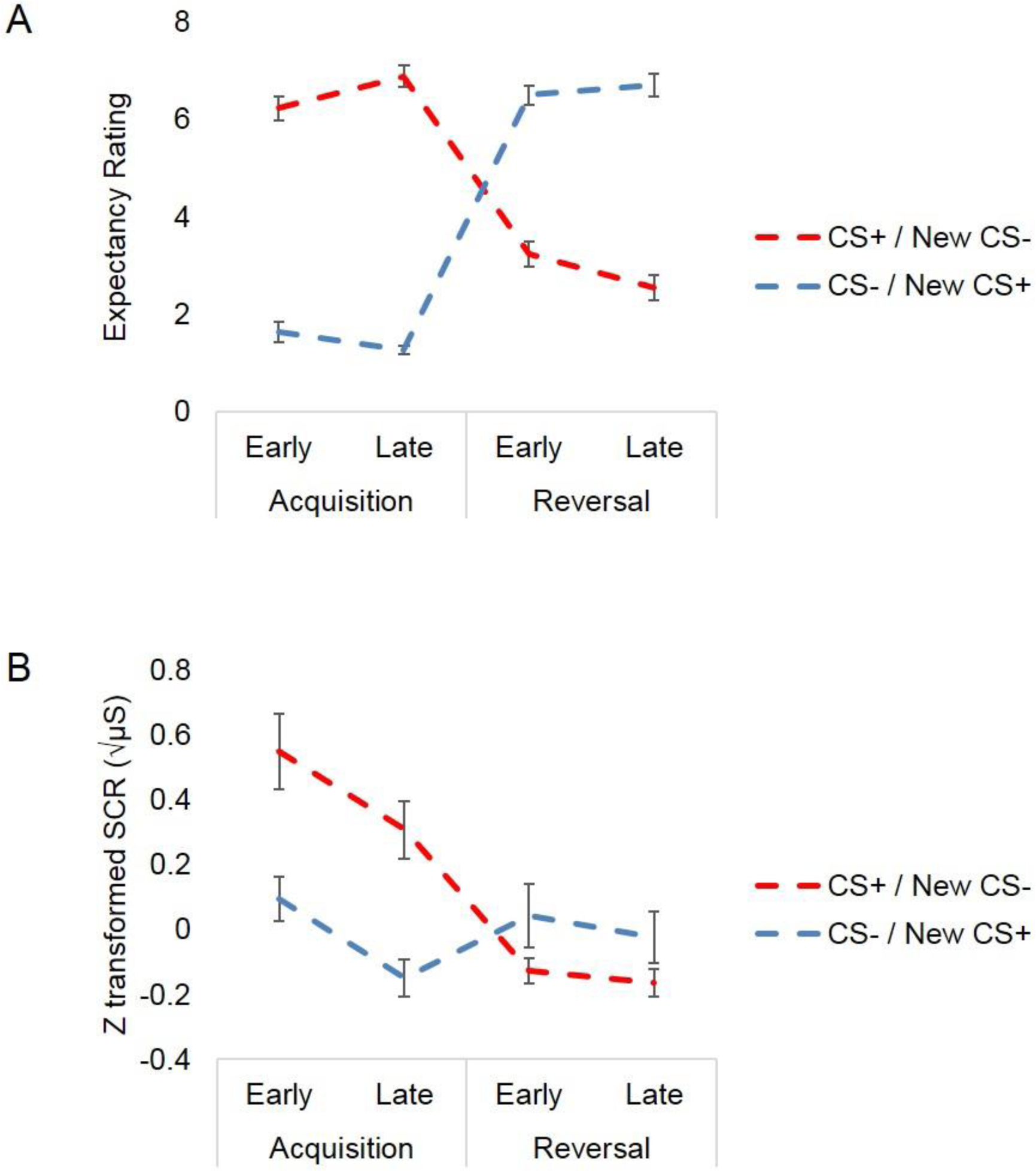
Line graphs displaying (A) expectancy ratings and (B) SCR magnitude scores to the CS+ and CS- during the experiment. For all phases participants reported greater expectancy of the sound with the CS+, compared to the CS-. In addition, larger SCR magnitude responses were found for the CS+ versus CS- during acquisition and reversal. Bars represent standard error of the mean. Expectancy rating, 1 ‘Don’t expect - 9 ‘Do expect’. SCR magnitude (√μS), square root transformed and z-scored skin conductance magnitude measured in microSiemens.

Reflecting learning over time, follow-up tests revealed the expectancy rating of the sound with the CS+ to increase from early acquisition to late acquisition, p < .05 [Stimulus x Time: *F*(1, 123.240) = 6.624, *p* = .011], and the expectancy rating of the sound with the CS- to remain low across early to late acquisition, *p* =.104. No other significant main effects or interactions with IU or STAIX-2 were found for the ratings during reversal, max *F* =.997.

During reversal, participants reported greater expectancy of the sound with the NewCS+, compared to NewCS- [Stimulus: *F*(1, 122.423) = 329.755, *p* < .001]. Follow-up pairwise comparisons suggest that the expectancy rating of the sound with the NewCS+ was high during this phase and did not change across early acquisition to late acquisition, p = .436 and the expectancy rating of the sound with the NewCS- dropped across early to late acquisition, *p* =.036 [Stimulus x Time: *F*(1, 122.423) = 4.691, *p* = .032]. No other significant main effects or interactions with IU or STAIX-2 were found for the ratings during reversal, max *F* = 2.289.

### 3.2 SCR magnitude

For SCR magnitude, during acquisition participants displayed larger SCR magnitude to the CS+, compared to CS- [Stimulus: *F*(1, 135.204) = 27.726, *p* < .001] (see Table 1 and Fig. 2b). Furthermore, SCR magnitude dropped from early acquisition to late acquisition without an effect of stimulus type [Time: *F*(1, 135.204) = 7.821, *p* = .006]. No other significant main effects were found, nor were there significant interactions with IU (or STAIX-2) for the SCR magnitude during acquisition, max *F* = .978.

During reversal participants displayed larger SCR magnitude to the NewCS+, compared to the NewCS- [Stimulus: *F*(1, 118.858) = 4.891, *p* = .029]. As predicted, MLM revealed a significant interaction between Stimulus x IU during reversal, *F*(1, 118.858) = 5.418, *p* = .022 (see Fig 3)^12^. Further inspection of follow-up pairwise comparisons for reversal revealed that low IU was associated with reduced and less differential responding between the NewCS+ vs. NewCS- during reversal, *p* = .353. High IU was associated greater differential responding to the NewCS+ vs NewCS- during reversal, *p* = .002. No other significant main effects or interactions with IU or STAIX-2 were found for the SCR magnitude during reversal, max *F* = 2.694.

**Fig 3.**
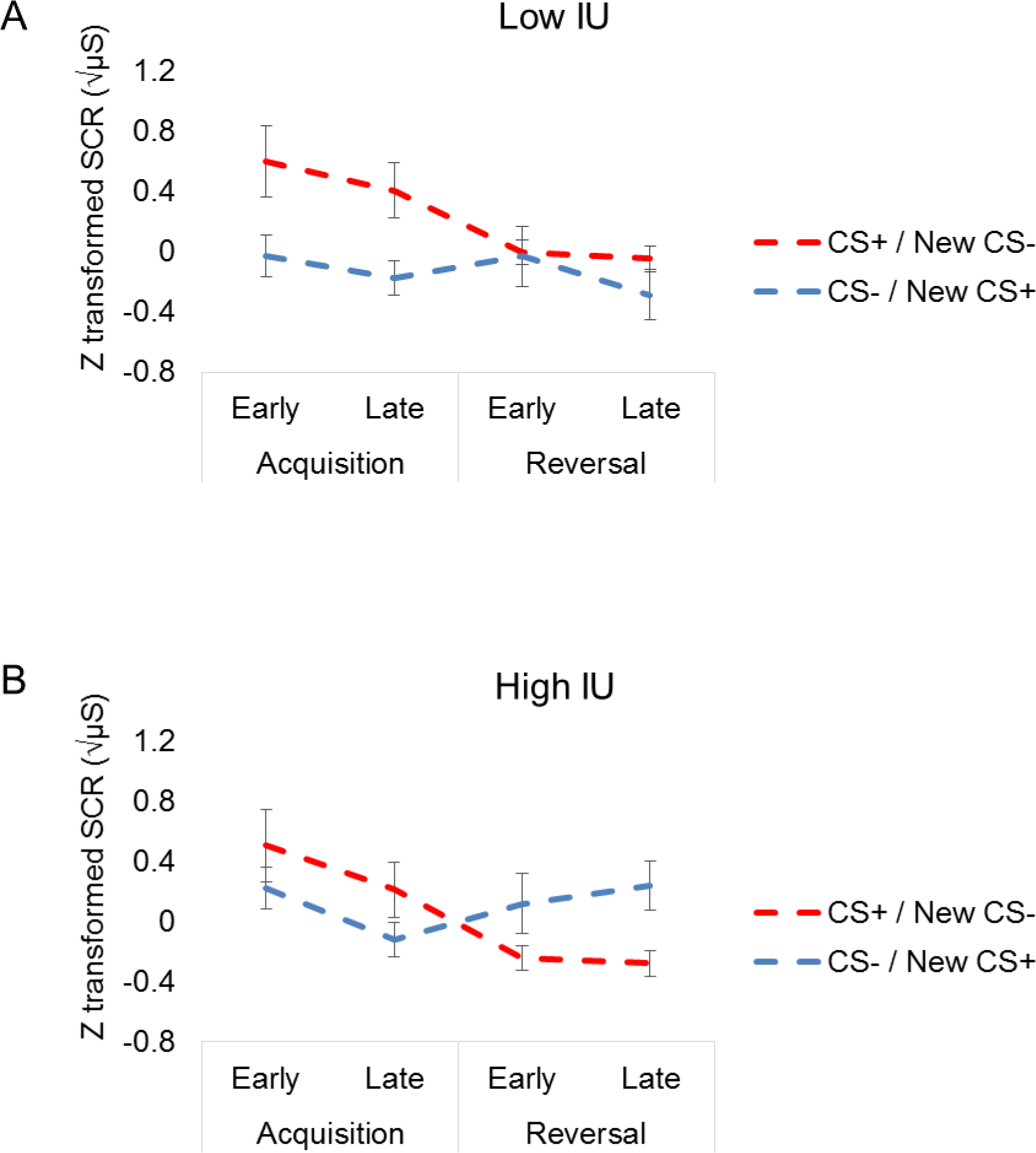
Line graphs depicting IU estimated at + or - 1 SD of mean IU (controlling for STAIX-2) from the multilevel model analysis for SCR magnitude during acquisition reversal. Higher IU was associated with greater discrimination in skin conductance responses to CS+ vs CS- during reversal. No significant IU-related differences were observed during acquisition. Bars represent standard error at + or – 1 SD of mean IU. Square root transformed and z-scored SCR magnitude (μS), skin conductance magnitude measured in microSiemens.

## 4. Discussion

In the current study, we showed that individual differences in self-reported IU modulate threat reversal. Our data suggest that during an associative learning experiment with acquisition and reversal phases, high IU is associated with greater differential skin conductance response between threat and safety cues during reversal. No significant relationships were found between IU and expectancy ratings. The skin conductance findings for IU are specific to variability explained by IU, over shared variability with STAIX-2. These results further our understanding of previous threat acquisition and extinction studies where psychophysiological and neural patterns of responding were associated with IU (Chin, Nelson, Jackson, & Hajcak, 2016; Dunsmoor, Campese, Ceceli, LeDoux, & Phelps, 2015; Morriss, Christakou, & Van Reekum, 2015, 2016; Morriss, Macdonald, & van Reekum, 2016). Furthermore, these results provide some preliminary findings on how threat reversal under partial reinforcement may be related to uncertainty-related biases, which may play an important role in the maintenance of uncertainty-induced anxiety.

For both threat acquisition and reversal phases, greater expectancy ratings and skin conductance magnitude were observed for threat versus safe cues, suggesting evidence of conditioning across the sample (Costa, Bradley, & Lang, 2015; Li, Schiller, Schoenbaum, Phelps, & Daw, 2011; Mertens & De Houwer, 2016; Morris & Dolan, 2004). However, the extent of conditioning on skin conductance magnitude during reversal varied substantially with individual differences in IU. During reversal, higher IU was associated with larger SCR magnitude to new threat vs. new safety cues, compared to lower IU. Interestingly, IU was not related to SCR magnitude in response to the US (see supplementary material). Taken together, these results suggest that IU-related biases may play a stronger role in modulating associative learning mechanisms rather than habituation based mechanisms.

In a broader context, the reversal finding with IU enriches our understanding of previous research where high IU has been found to predict reduced threat extinction (Morriss et al., 2015; Morriss, Christakou, et al., 2016; Morriss, Macdonald, et al., 2016). The current results suggest that individuals with high IU scores are able to update threat and safety associations in the presence of threat. The reason for high IU individuals being able to update threat to safe associations during reversal but not extinction may stem from the amount of information available. For example, there is some familiarity with the reinforcement rate, such that CSs are associated with a 50% partial reinforcement rate during both acquisition and reversal. Furthermore, during reversal, the learned threat is simply moved onto another stimulus, whilst in extinction, the learned threat is completely omitted. The lack of any previously learned threat outcome in extinction may induce greater anxiety in high IU individuals. This finding would fit with the modern definition of IU, ‘IU is an individual’s dispositional incapacity to endure the aversive response triggered by the perceived absence of salient, key, or sufficient information, and sustained by the associated perception of uncertainty’ (Carleton, 2016b, p. 31). Further work is needed to clarify how the presence and absence of information modulates anxiety in individuals who score high in IU.

The findings reported here feed into a wider research movement examining the role of uncertainty in anxiety disorders (Carleton, 2016a, 2016b; Grupe & Nitschke, 2013). Recent research has begun to examine whether IU can be targeted in treatment for anxiety disorders and initial findings show promise for patients with generalised anxiety disorder and social anxiety disorder (Dugas & Ladouceur, 2000; van der Heiden, Muris, & van der Molen, 2012). Our preliminary findings highlight how IU interacts with fundamental associative learning processes such as threat conditioning, which are relevant for current exposure-based therapies for anxiety disorders (Milad & Quirk, 2012). It is unclear, however, what type of IU-related biases interact with threat conditioning mechanisms e.g. one potential bias could be heightened expectation of threat under uncertainty, particularly when direct threat is absent as in extinction. Nonetheless, such findings pave the way for new lines of research to address the relevance of IU-related biases on threat conditioning mechanisms as potential risk markers and intervention targets. For example, future research may focus on how discrimination during threat conditioning can be improved in high IU individuals, by exposing participants to more safe stimuli or by asking participants to use different appraisal strategies.

In the current study, we did not observe IU to modulate threat acquisition. This result is at odds with some previous research from our lab and others, where high IU individuals has been shown to have: (1) larger SCR magnitude across threat and safety cues during partial reinforcement (Morriss, Macdonald, & van Reekum, 2016) and (2) have greater discrimination in startle response for threat versus safe cues (Chin, Nelson, Jackson & Hajcak, 2016). The differences in findings may reflect differences in experimental paradigms and measures. For example, there may be other differences between the studies which may influence the results, such as the threat level of unconditioned stimuli (e.g. sound versus shock), number of stimulus types (e.g. the use of generalisation stimuli) and the number of trials. Furthermore, startle is under control of the central nucleus of the amygdala, among other (Koch, 1999), and the inclusion of startle probes introduces more ambiguity in the experimental design (Lonsdorf & Merz, 2017). Further research is needed to assess the role of IU in threat acquisition.

Self-reported expectancy ratings were not found to reflect individual differences in IU in our sample. Whilst correspondence between ratings and SCR magnitude was observed, differences between self-reported and psychophysiological measures of emotion are often reported (Mauss, Levenson, McCarter, Wilhelm, & Gross, 2005). To our knowledge only a few studies have observed IU effects on ratings during threat conditioning (Morriss, Macdonald, & van Reekum, 2016; Sjouwerman, Scharfenort, & Lonsdorf, 2017). The majority of research examining the effects of IU on threat acquisition and extinction have found significant relationships between IU and psychophysiological measures such as startle and skin conductance (Chin, Nelson, Jackson, & Hajcak, 2016; Dunsmoor, Campese, Ceceli, LeDoux, & Phelps, 2015; Morriss, Christakou, & Van Reekum, 2015, 2016; Morriss, Macdonald, & van Reekum, 2016; Sjouwerman, Scharfenort, & Lonsdorf, 2017). We therefore think that IU may be a more suitable predictor of bodily responses during threat conditioning, potentially capturing both unconscious and conscious processing. The lack of relationship between psychophysiological and rating measures for IU may also be due to the time between phasic cue events and rating periods in the experiment, where recall of expectancy was required for each block.

The design specifics of the current study should be further addressed in future research to assess the robustness and generalizability of the findings reported here. Firstly, the generality of these findings should be tested in future studies using stimuli other than coloured squares and sounds. Secondly, using different reinforcement rates – ideally in a single study - may elucidate if individuals high in IU are more sensitive to some reinforcement rates over others during acquisition and reversal. Thirdly, the sample contains mainly female participants, and future studies should more carefully balance their sample in terms of gender. Lastly, the current study may have been more sensitive to individual differences if low and high IU individuals were selected or over-sampled. All these points would benefit from further research to assess the robustness and generalizability of the findings reported here.

In conclusion, these initial results provide some insight into how threat reversal under partial reinforcement may be related to IU, which may be relevant for the understanding of uncertainty-induced anxiety (Carleton, 2016a, 2016b; Grupe & Nitschke, 2013). Further research is needed to explore how individual differences in IU modulate learned threat and safety associations.

## Acknowledgements

This research was supported by the Centre for Integrative Neuroscience and Neurodynamics (CINN) at the University of Reading. The authors thank the participants who took part in this study. To access the data, please contact Dr. Jayne Morriss.

## Supplementary material: US habituation

To examine whether US habituation to the CS+paired trials was related to IU and STAIX-2, we conducted additional analyses.

### Physiological scoring of US responses to CS+ paired trials

The amplitude of each SCR was scored as the difference between the onset and the maximum deflection prior to the signal flattening out or decreasing. SCR onsets and respective peaks were counted if the SCR onset was within 4.5-8 seconds (US response) following CS+ paired onset. Trials with no discernible SCRs were scored as zero. SCR amplitudes were square root transformed to reduce skewness. SCR magnitudes were calculated from remaining trials by averaging SCR square root transformed values and zeros for each contingency and block, creating the following conditions; Acquisition CS+paired Early (N=3); Acquisition CS+paired Late (N=3); Reversal New CS+paired Early (N=4); Reversal New CS+paired Late (N=4). SCR magnitudes were z-scored across CS+paired trials to control for interindividual differences in skin conductance responsiveness related to dryness and thickness of the skin and the surface area of the finger relative to the skin conductance electrode.

### SCR magnitude analysis of US responses to CS+ paired trials

The analysis was conducted using the mixed procedure in SPSS 21.0 (SPSS, Inc; Chicago, Illinois). For SCR magnitude we analysed each phase separately, we entered Time (Early, Late) at level 1 and individual subjects at level 2. We included the following individual difference predictor variables into the multilevel models: IU, STAIX-2. In all models, we used a diagonal covariance matrix for level 1. Random effects included a random intercept for each individual subject, where a variance components covariance structure was used. Fixed effects included Time. We used a maximum likelihood estimator for the multilevel models.

### Results of US responses to CS+ paired trials

For both acquisition and reversal, no significant main effects of Time or interaction with IU or STAIX-2 were observed, max *F* = 1.753.

### Footnotes

1 In the untransformed SCR data, the same pattern of results are observed for the interaction of Stimulus x IU during reversal, *F*(1,95.368) = 5.760, *p* = .018.

2 To assess that the interaction of Stimulus x IU during reversal was not driven by US habituation, we ran an additional MLM including the average SCR magnitude of the US response during reversal as a covariate. In this model, we observed the same result for Stimulus x IU during reversal, *F*(1,95.368) = 4.757, *p* = .031, suggesting that this effect was not driven by US habituation.

---

**Table S1.**
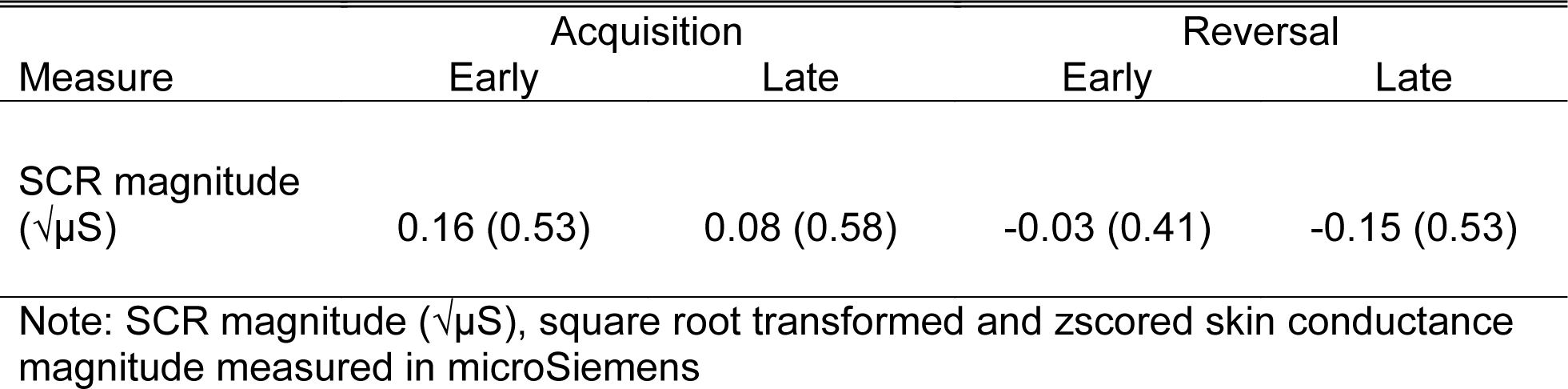
Summary of means (SD) for SCR magnitude to CS+paired trials during the acquisition and reversal phases

